# Evolution and expression of Glial Cells Missing (*GCM1* and *GCM2*) in monotremes suggests an ancient role in reproduction and placentation

**DOI:** 10.1101/2025.03.16.643584

**Authors:** Isabella Wilson, Diana Demiyah Mohd Hamdan, Frank Grützner

## Abstract

The Glial Cells Missing (GCM) genes were first discovered in *Drosophila* and encode transcription factors important for gliogenesis. In placental mammals, *GCM1* regulates several genes that are important for early placenta development, while its paralog *GCM2* is important for parathyroid gland development. The egg-laying monotremes, which represent the most diverged extant mammalian lineage, undergo a short period of intrauterine development and form a simple placenta. To gain more insight into the evolution of *GCM* genes we analysed the sequence, expression, and genomic localization of *GCM1* and *GCM2* genes in the platypus and echidna. We found that the chromosomal localisation of *GCM1* changed after the divergence of therian mammals, coinciding with the evolution of a complex placenta. Expression analysis revealed the presence of GCM transcripts in male and female monotreme gonads, as well as expression of *GCM1* in the female reproductive tract. GCM binding sites in target genes associated with placental development in therian mammals were also present in the monotremes and the chicken. Together, this suggests that the role of *GCM1* in the placenta emerged early in mammalian evolution.

## Background

The glial cells missing (GCM) transcription factors are a small, highly evolutionarily conserved family of genes involved in development. In *Drosophila, gcm* is involved in cell differentiation during early neurogenesis^1^. Amniote genomes contain two GCM genes; *GCM1* is required for placenta development whereas *GCM2* plays a role in the development of the parathyroid^2^.

The placenta is an autonomous, short-lived organ which serves the essential function of supporting the development of offspring during intrauterine gestation^3^ *GCM1* is master regulator of placental cell differentiation, playing key roles in facilitating placental invasion and maternal-fetal nutrient exchange. In the human and mouse placenta, *GCM1* is expressed in a subset of cytotrophoblasts where it governs syncytiotrophoblast differentiation and villous morphogenesis^4,5^. In humans, alterations to *GCM1* gene expression are associated with severe placental insufficiency syndromes such as preeclampsia and intrauterine growth restriction^6^, and *gcm1* knockout in mice is embryonic lethal^7^. In addition to its expression in placenta, low levels of *gcm1* transcript are also detected in adult kidney tissues^8^.

As one of the master regulators in placentation, *Gcm1* is required to up-regulate the expression of a range of target genes in the placental mammals^9,10^. The N-terminal region of the GCM proteins in vertebrates consists of a highly evolutionary conserved DNA-binding domain, which binds as a monomer to its DNA binding sites in a sequence-specific manner. Using a β-sheet, the GCM DNA-binding domain can specifically recognise an octameric sequence motif in potential target genes^11–13^. Two key targets of *GCM1* are the syncytin genes *ERVW-1* and *ERVFRD-1*, retroviral genes that have fusogenic properties mediating cell-cell fusion between trophoblasts to form the multinucleate syncytial layer of the placenta^14–17^. *GCM1* upregulates the expression of the syncytins and their receptor *MFSD2A* in the placental syncytiotrophoblast^18,19^. *GCM1* is also important in placental vasculogenesis, playing a key role in regulating *PGF* expression in trophoblasts^9,20^. *Gcm1*-deficient mice exhibit reduced expression of genes involved in labyrinthine layer formation (*Itga4* and *Rb1*), suggesting that *GCM1* is important in facilitating maternal-fetal nutrient exchange^10^.

The transcriptional activity of GCM1 appears to be highly conserved throughout evolution, with localisation of expression playing a role in its changing function. Expression of recombinant murine *GCM1* in mouse embryonic brain cells has been demonstrated to induce glial differentiation^21^. Furthermore, when murine *Gcm1* is expressed in the developing nervous system of *Drosophila*, extra glial cells are generated at the expense of neurons^22^. This indicates evolutionary conservation of the regulatory capabilities of *GCM1*. In chicken, *Gcm1* is transiently expressed during embryogenesis in the chorioallantoic membrane^23^. The localisation of *Gcm1* expression in tissues that facilitate nutrient supply to the embryo in both chicken and placental mammals suggests an evolutionarily conserved role in early foetal development.

Deep evolutionary divergence and unique reproductive biology (egg-laying and simple placenta) make monotremes (platypus and echidna) a key species in which to investigate genes involved in placentation. Though they are oviparous, monotremes develop a simple yolk sac placenta which supports the embryo during a short intrauterine development (∼21 days in platypus, ∼23 days in echidna^24,25^). Being oviparous, monotreme placentae do not make direct contact with the uterine epithelium and are thus non-invasive. However, the thin shell coat is permeable such that physiological exchange is possible, and the placenta is known to undergo a period of expansion to maximise its surface area (and therefore its potential for nutrient exchange)^26^. The thin trilaminar trophectoderm that forms could theoretically be capable of haemotrophic exchange, though there are no studies to verify this^27^. At the molecular level, monotreme placentation is still poorly understood. Many of the genes associated with maternal-foetal immunotolerance were recruited into endometrial expression after the divergence of therian mammals, a process understood to be orchestrated by transposable elements^28^. The retrotransposon-derived gene *PEG10,* which is important for placenta formation in mouse, is missing in monotremes^29^. *POU5F1*, a gene that is essential for trophectoderm differentiation^30,31^ has been shown to be conserved and functional in the platypus^32^.

In order to gain a better understanding of placental evolution and function, we characterised the GCM genes in the platypus and short-beaked echidna and investigated the conservation of the binding sequence of *GCM1* target genes between amniotes. This revealed that monotremes exhibit expression of GCM genes in reproductive tissues, and that GCM binding sites within placenta-associated target genes are conserved even in species that lack a complex placenta, suggesting that the reproductive function of *GCM1* emerged early in mammalian evolution.

## Results

### Characterisation of platypus and echidna GCM orthologues reveals changes after divergence of monotremes

GCM orthologs were identified in the most recent annotation releases for the platypus and echidna genomes^33,34^. In both the platypus and the echidna, *GCM1* is annotated on chromosome 1 and *GCM2* is annotated on chromosome X2. We sought to confirm the physical location of *GCM1* and *GCM2* using fluorescence in-situ hybridisation (FISH) on platypus tissue. As expected, we observed a signal for *GCM1* on chromosome 1 (Fig 1A).

**Figure 1:**
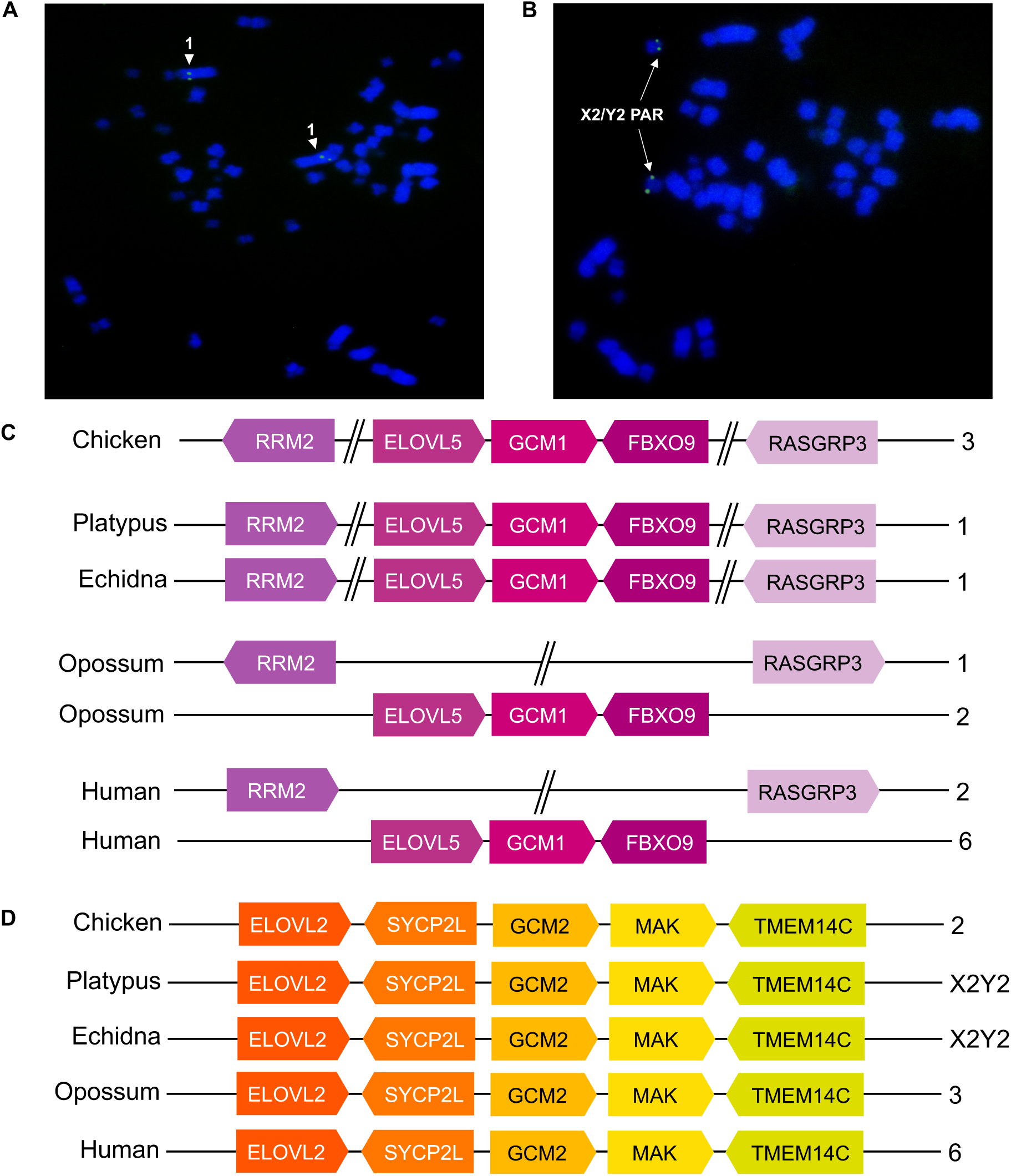
Chromosomal localisation of GCM genes. Fluorescence *in situ* hybridization of BAC clones (A) OABb-307B16 (green) and (B) OABb-373O17 (green) on male platypus metaphase chromosome spreads. Chromosomes were counterstained with DAPI. (C) Schematic showing the chromosomal location of *GCM1* orthologues and neighbouring genes in the platypus, echidna, human, opossum, and chicken. (D) Syntenic arrangement of *GCM2* in amniotes.

Physical mapping of *GCM2* showed localisation to the X2Y2 pseudoautosomal region (Fig. 1B). We attempted to locate the Y2 copy of *GCM1* by performing a tBLASTn search against the translated platypus and echidna genomes using both the full GCM2 protein sequence and the GCM2 DNA-binding domain, but no matches were retrieved.

Next, we investigated the genomic context of GCM genes in monotremes. This showed conservation of synteny of *GCM1/2* flanking genes across 5 species: chicken, platypus, echidna, opossum, and human. In the platypus, echidna, and chicken, genes neighbouring *GCM1* included *RRM1*, *ELOVL5*, *FBXO9*, and *RASGRP3* (Fig. 1C). However, synteny breaks occurred in therian mammals involving adjacent genes *RRM2* and *RASGRP3* (Fig. 1C). For *GCM2*, we found that neighbouring genes (*ELOVL2*, *SYCP2L*, *MAK*, and *TMEM14C*) had the same syntenic arrangement as in other amniotes, showing homology between platypus and echidna chromosome X2, human chromosome 6, opossum chromosome 3 and chicken chromosome 2 (Fig. 1D).

### Sequence-level characterisation of monotreme *GCM1* and *GCM2*

A phylogenetic tree was constructed to analyse the conservation of the monotreme GCM sequences in comparison to other vertebrates (Supplementary Figure 1). In general, the topology of the tree was as expected. The functional domains of the monotreme sequences were analysed in order to better understand potential functional changes as a result of amino acid substitutions (Figure 2). Analysis of the DNA-binding domain (termed the GCM domain) in the platypus and echidna confirmed high conservation for both *GCM1* and *GCM2*. The GCM domain is stabilised by two Zn ions which bind to a series of conserved cysteine and histidine residues of the sequence C-X48-C-X26-H-X-H and C-X3-C-X26-C-X2-C^13^. This cysteine and histidine pattern was conserved in the monotremes. Furthermore, all DNA-contacting residues within the GCM domain were conserved between monotremes and other amniotes for both GCM amino acid sequences, with the exception of one non-conservative replacement at position 102 unique to the echidna GCM1 sequence. This suggests that monotreme GCM proteins are able to bind target DNA. However, monotreme-specific changes were observed in the GCM1 transactivation domains (Fig. 2A). These included a 9aa deletion in transactivation domain 1 and two 3aa insertions in transactivation domain 2, which could affect the ability of monotreme GCM transcription factors to activate target genes.

**Figure 2:**
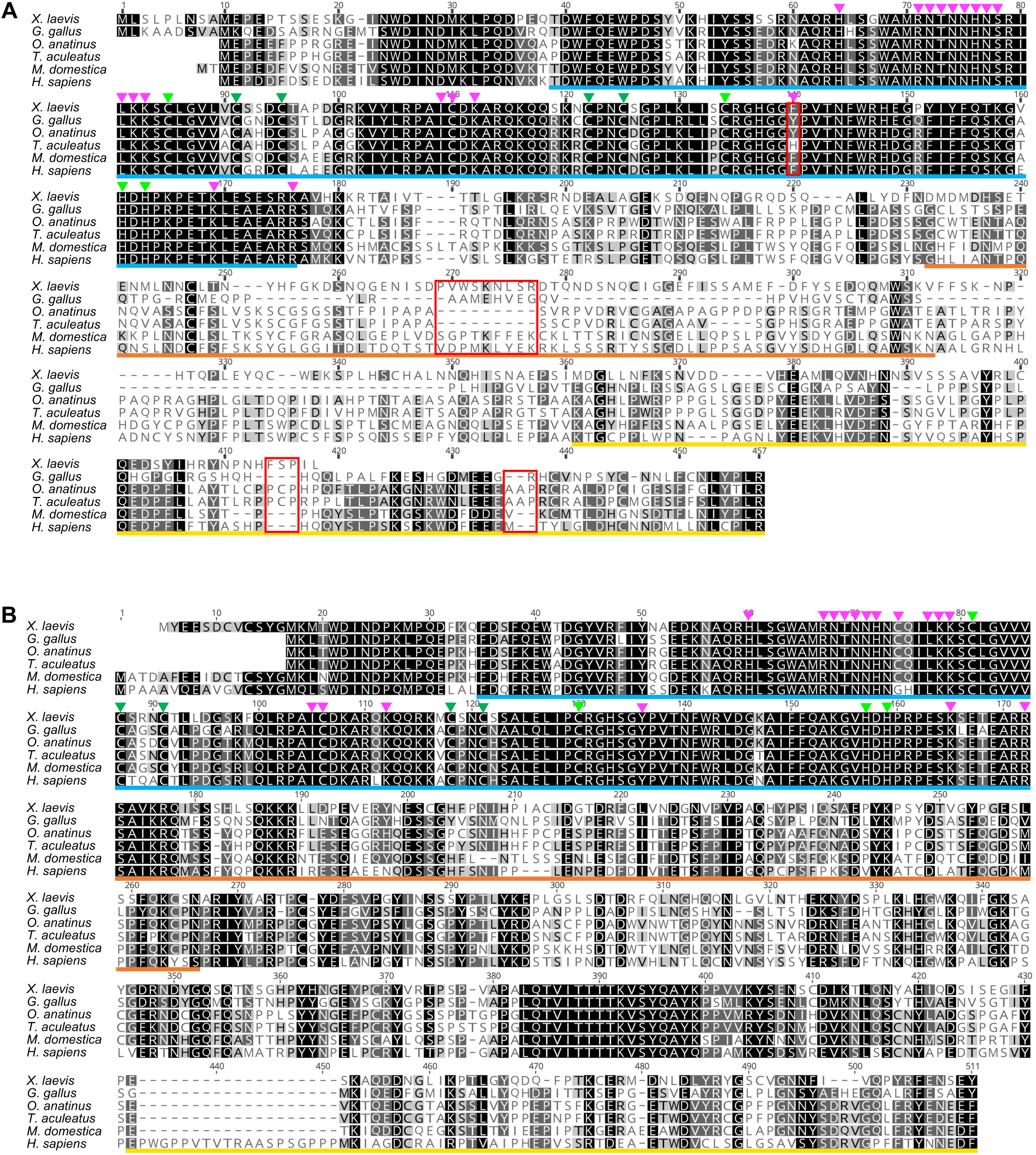
Conservation of GCM1 (A) and GCM2 (B) in monotremes and other vertebrates. GCM DNA binding domain is underlined in blue. Light and dark green arrows indicate residues coordinating first and second Zn2+ respectively. Magenta arrows indicate DNA-contacting residues. Transactivation domain 1 is underlined in orange. Transactivation domain 2 is underlined in yellow. Notable sequence changes are indicated by red boxes. Shading indicates similarity of amino acid residues.

### GCM expression in adult monotreme tissues

Expression of *GCM1* and *GCM2* was investigated by RT-PCR in adult male and female platypus and echidna tissues. *GCM1* was expressed in the testis and female reproductive tract of both monotremes (Fig. 3A, 3B). Female reproductive tract expression of *GCM1* seemed stronger in echidna than in platypus. *GCM1* and *GCM2* were strongly expressed in platypus ovary (Fig. 3A), though surprisingly no expression was observed in echidna ovary (Fig. 3B). *GCM1* transcript was also detected in platypus and echidna kidney tissues in both males and females. In placental mammals, expression of *GCM1* in gonads has not been reported and we did not detect any expression of *Gcm1* in adult mouse gonads (Fig. 3C). Our experiments confirm expression of *Gcm1* in adult mouse kidney^8^.

**Figure 3:**
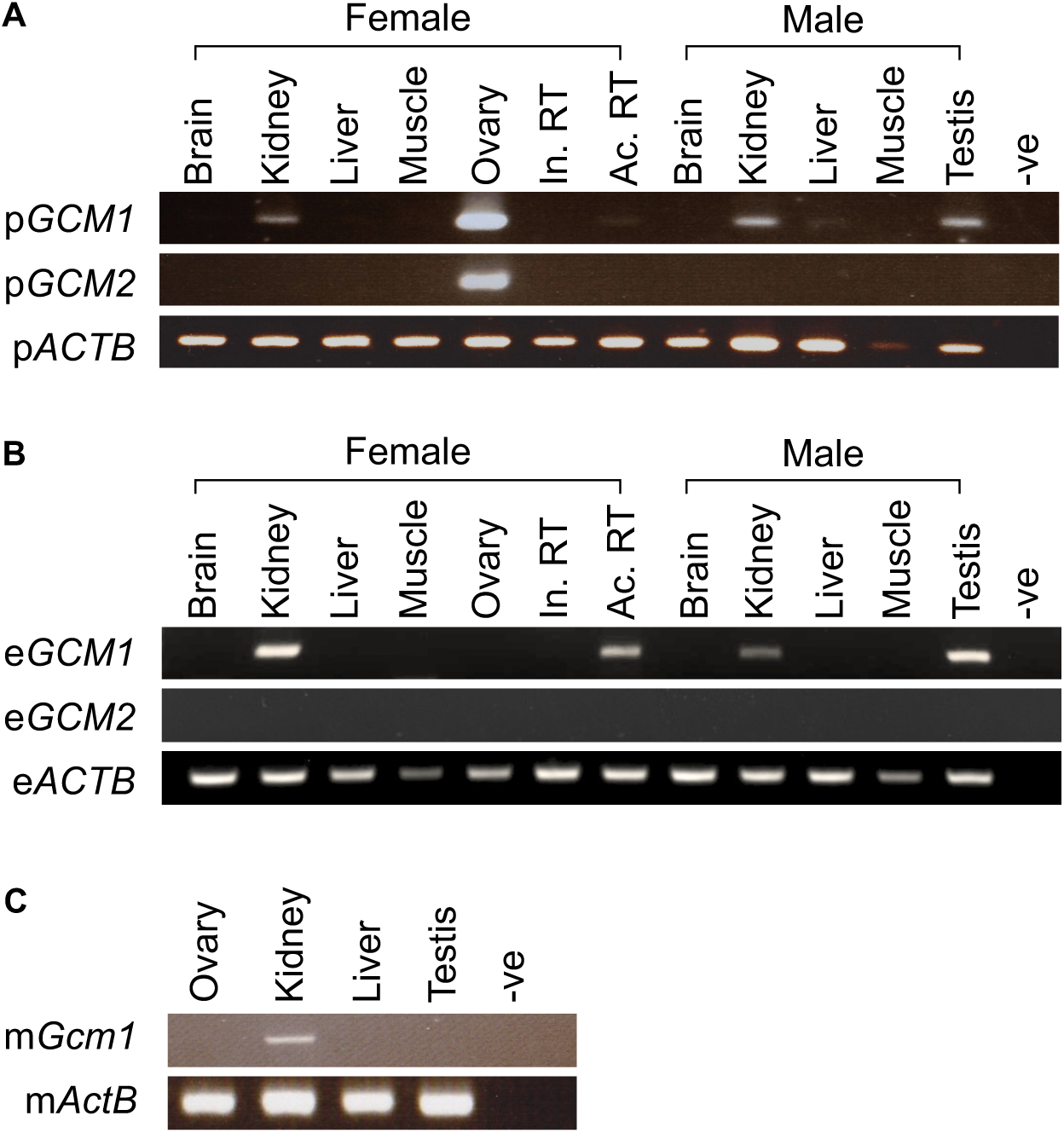
RT-PCR showing expression of *Gcm1* and *Gcm2* in platypus and echidna tissues. PCR experiments were performed using cDNA from adult male and female tissues from (A) platypus, (B) echidna, and (C) mouse. An *ACTB* PCR was performed on all tissues as a positive control. A negative control (no template) was included for each PCR experiment. Abbreviations: In. RT = inactive reproductive tract, Ac. RT = active reproductive tract, -ve = negative control.

### Expression pattern of *GCM1* and *GCM2* in platypus reproductive tissues

In order to investigate the expression pattern in platypus gonads in more detail, we conducted RNA *in situ* hybridization of *GCM1* and *GCM2.* In platypus adult testis (samples taken during breeding season) in three different individuals, expression of *GCM1* is detected in the germ cells in the seminiferous tubules (Fig. 4A). Platypus *GCM1* transcript was not detected in the interstitial connective tissue of the testis (Fig. 4A). Platypus *GCM2* expression was detected in adult ovary, showing expression in granulosa cells and interstitial cells (Fig 4D). We observed co-expression of platypus *GCM1* and *GCM2* in ovary (Fig. 4C, 4D). In the platypus female reproductive active tract, expression of *GCM1* was restricted to a specific subset of cells in the myometrium (Fig. 4B).

**Figure 4.**
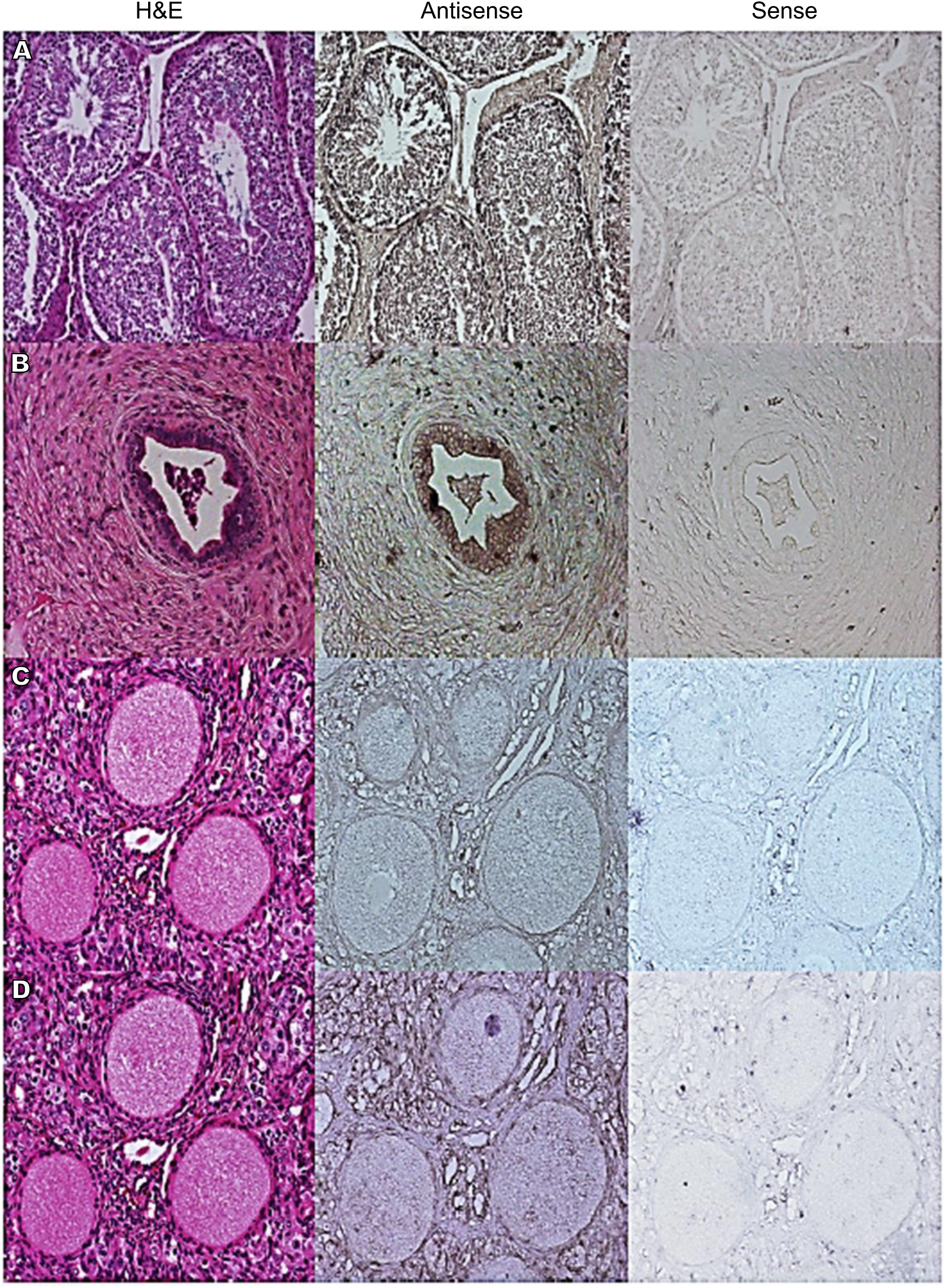
Detection of platypus *GCM1* and *GCM2* transcripts in reproductive tissues. (A) *GCM1* was detected in the germ cells of the seminiferous tubules in testis (X10 magnification). (B) *GCM1* expression was restricted to specific subsets of cells in the female reproductive tract myometrium (X20 magnification). (C) Expression of *GCM1* was detected in ovary interstitial cells (X20 magnification). (D) *GCM2* was expressed in ovary interstitial cells (X20 magnification). Left column shows tissue sections stained with H&E. Middle column shows tissue sections hybridised with antisense probe. Right column shows tissue sections hybridised with sense probe as a control.

### Conservation of GCM DNA binding sites within target genes in amniotes

GCM1 upregulates the expression of its target genes by binding to GCM DNA-binding sites^9,19^. The GCM domain recognises DNA-binding sites with the motif sequence 5’-(A/G)CCC(T/G)CAT-3’or its compliment 5’-ATG(A/C)GGG(T/C)-3 in target genes^11,12^. We identified monotreme orthologues of known GCM target genes expressed in the placenta, including *ITGA4*, *MFSD2A*, *RB1*, *PGF*, and *PITX2*.

In order to identify candidate GCM binding sites, the target motif sequence was identified within target genes in addition to the 1000 bp immediately preceding the transcription start site. This revealed that in both the platypus and the echidna, GCM DNA-binding sites are present in GCM target gene orthologues (detailed sequence information can be found in Supplementary Data File 1). Surprisingly, the target motif sequence was also present in chicken orthologues of *ITGA4*, *MFSD2A* and *RB1* despite these genes lacking a placenta-associated role in birds.

In order to analyse whether individual target sites were conserved across species, we aligned target gene orthologues (plus 1000 bp upstream of the translation start site) and compared the position and sequence of each putative GCM DNA-binding site (Figure 5). The two monotremes showed a higher level of similarity to each other than to other species, presumably due to their close evolutionary distance. However, in general, the location of binding sites varied greatly along orthologous gene sequences, suggesting that although each target gene orthologue possesses GCM DNA-binding sites, specific sites are not conserved. One exception was a binding site located across an intron/exon boundary in *ITGA4* which was present in all species investigated.

**Figure 5:**
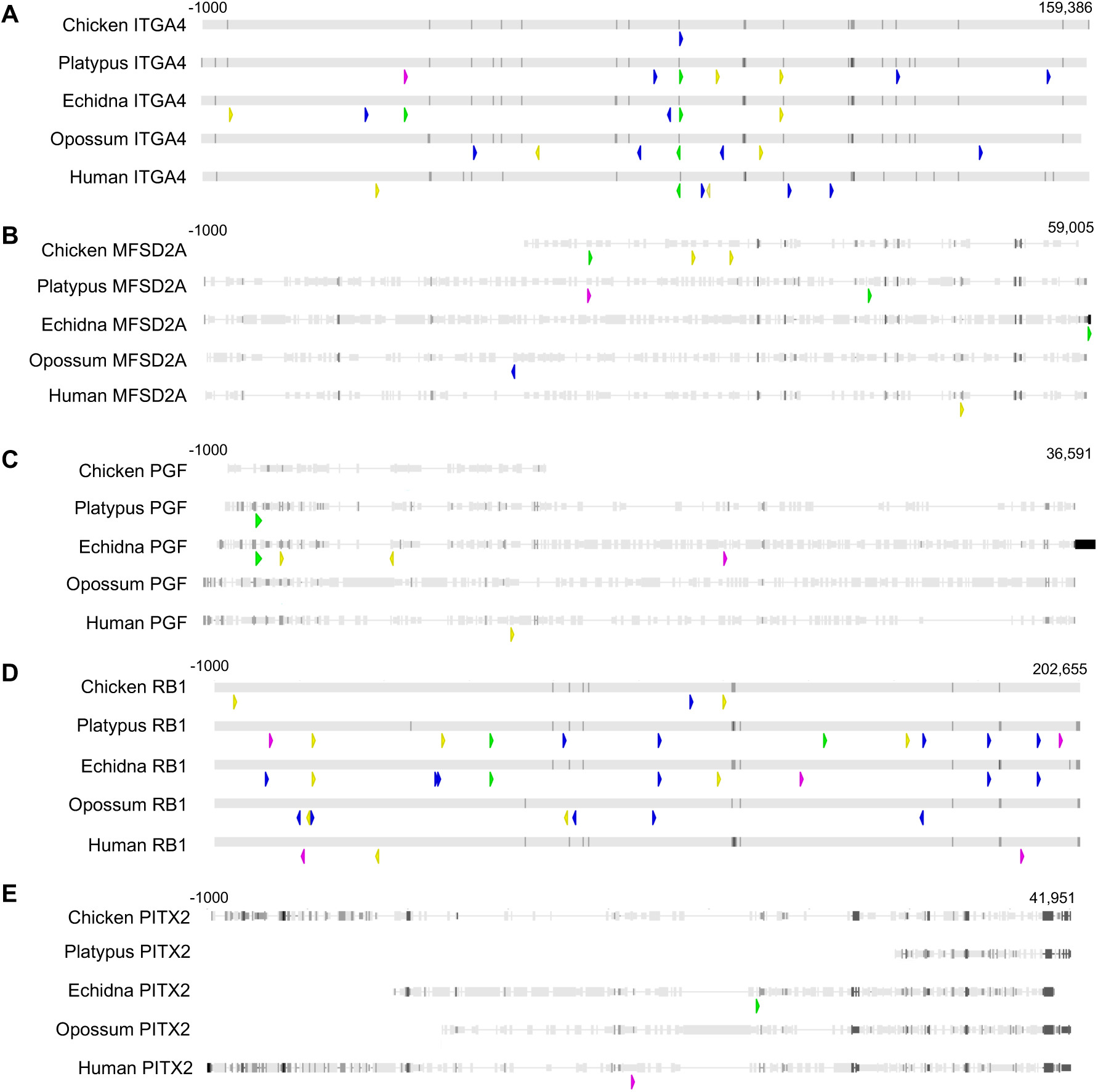
Conservation of GCM binding site motifs in orthologues of GCM target genes in amniotes. DNA binding sites are indicated with arrows; each colour corresponds to a particular motif sequence (blue = ACCCTCAT, yellow = GCCCTCAT, green = ACCCGCAT, pink = GCCCGCAT).

GCM1 is also known to regulate the syncytin gene *ERVW-1* in humans. A recent study revealed over 120 putative syncytin genes in the echidna genome^35^. Human *ERVW-1* and echidna syncytin genes are not orthologous as they are retroviral and inserted into the different lineages independently, however we were curious to see whether the putative echidna syncytins contained GCM DNA-binding motifs. After searching the putative syncytin ORFs for the GCM target motif sequence, we discovered 59 GCM target sites, with 55 out of a total 121 putative syncytin ORFs containing at least one GCM target motif (Supplementary data file).

## Discussion

The evolution of a complex placenta is a hallmark of mammalian evolution. All mammalian lineages feature a placenta, including the egg-laying monotremes, which develop a simple yolk-sac placenta during a short period of intrauterine development^36^. Though mammalian placentae share a basic nutrient-waste exchange function, major differences in structure and invasiveness can be observed between lineages. Here we present the first analysis of an essential placental development gene, *GCM1,* in the platypus and echidna.

Chromosome analysis revealed that a syntenic block containing *GCM1* and its neighbouring genes has been largely conserved in amniotes but was relocated to a different chromosome after the divergence of therian mammals (Fig 1C). *GCM2* also underwent changes with respect to its chromosomal localisation throughout mammalian evolution; monotreme *GCM2* is located on a sex chromosome (Fig 1D). It is unclear whether the localisation of *GCM2* on sex chromosomes has any relevance in terms of function and evolution in the platypus ovary and absence of expression in echidna ovary. Our physical mapping demonstrates *GCM1* copies on both X2 and Y2 (Fig 1B). Together with the platypus sex chromosome pseudoautosomal boundaries recently defined by Zhou et. al^33^, this suggests that *GCM2* resides within the X2/Y2 pseudoautosomal region. However, attempts to identify the Y2 copy of *GCM2* in the platypus and echidna genomes using computational approaches were unsuccessful.

The amino acid sequence of the GCM domain exhibited a high level of conservation between birds, monotremes, and therian mammals (Fig 2). This suggests that platypus and echidna GCM proteins are able to bind target genes in much the same way as is seen in other species. However, monotreme-specific changes were observed in transactivation domains 1 and 2 of *GCM1*. These domains are associated with stability and transcriptional activity and in mice are thought to interact with the DNA binding domain^13^. The effects of small changes to transactivation domains are not well understood, but it is possible that the observed deletions in the monotreme *GCM1* transactivation domains could impact its transcriptional activity.

We observed *GCM1* expression in the active reproductive tract of both the platypus and the echidna (Fig 3). Despite their oviparity, the reproductive tract plays a role in maternal-foetal physiological exchange in the monotremes by producing nutrient secretions which are able to pass through the permeable shell coat and across foetal membranes^37^. The expression of *GCM1* in the monotreme reproductive tract suggests an ancient role in maternal-fetal nutrient exchange, which predates the evolution of viviparity and complex invasive placentae. The absence of *GCM1* expression in the inactive reproductive tract further suggests that the role of *GCM1* in this tissue is associated with pregnancy and potentially with nutrient exchange. This conserved role is supported by research showing expression of *GCM1* in the chorioallantoic membrane of chicken embryos^23^, suggesting a role for this gene in maternal-fetal exchange even before the divergence of mammals.

In most vertebrates, *GCM2* expression is restricted to the pharyngeal pouch and the parathyroid gland where it acts as a master gene regulator in parathyroid gland development^2,38,39^. Lack of material prevented us from investigating GCM expression in platypus and echidna parathyroid tissue. However, we observed *GCM2* expression in platypus adult ovary (Fig. 3A). Ovarian expression has so far not been reported in any other vertebrate species. *GCM2* expression is typically associated with parathyroid cells, a hormone-producing cell-type of the parathyroid gland^40^. In the platypus, expression of *GCM2* was observed in granulosa and interstitial cells, both of which are associated with hormone production in the ovary. *GCM2* was not expressed in other platypus tissues tested (brain, kidney, liver, muscle, testis, reproductive tract, and echidna ovary), or in any echidna tissues. The lack of expression in echidna ovary is surprising given the high level of sequence similarity between the platypus and echidna *GCM2* sequences (Figure 2).

One explanation for the unique expression of *GCM2* in the platypus ovary is that the differences between platypus and echidna ovarian development and function could be responsible for divergent *GCM2* expression. One of the marked differences between ovarian development in the platypus as compared to echidnas, marsupials and most placental mammals is that only one ovary is fully functional^41^. This is also observed in chickens, where during early female chicken embryogenesis, asymmetrical ovary development results in only one functional ovary^42^. In chickens, this asymmetric development is controlled by the homeodomain transcription factor *Pitx2*^43^, which in mice is co-expressed with *Gcm1* in the placenta and kidney^44^. *Pitx2* also influences gonadal development in mammals; different isoforms of rat *Pitx2* exhibit sexually dimorphic expression during embryonic and postnatal ovary and testis development^45^. It is possible that the unique expression of GCM genes in the platypus ovary is related to asymmetric gonad development. We observed co-expression of *GCM1* and *GCM2* in the interstitial cells and granulosa cells of adult ovary tissue rather than in theca cells as is reported for rat *Pitx2* (Fig. 4C, 4D). The platypus *GCM1* transcript expression in testis is restricted to the seminiferous tubules and germ cells (Fig. 4A), resembling the localisation of rat *Pitx2* transcript detected in postnatal rat testis tissue^45^. Our analysis of GCM binding sites in monotreme *PITX2* orthologues suggests that there is one binding site in the echidna and none in the platypus (Fig 5E); however, the existing platypus *PITX2* gene annotation appears truncated in comparison to other amniote orthologues and may not be representative of the true platypus *PITX2* sequence. The co-expression and/or interaction of GCM genes with *PITX2* in platypus gonadal tissue remains to be demonstrated. Alternatively, the difference in ovarian GCM expression between platypus and echidna could be related to the samples themselves. While samples were taken from pregnant individuals for both platypus and echidna, it is unknown whether the animals were at different stages of gestation. Research in mice and chickens has demonstrated that GCM1 is expressed during specific stages of embryogenesis^46–48^. It is possible that monotreme GCM expression in ovary is also timed; further samples need to be investigated to confirm this.

A number of genes regulated by *GCM1* have been identified in placental mammals^9,18,19^. Members of the GCM family of transcription factors bind to target DNA in a sequence-specific manner; the optimum sequence bound by the GCM DNA-binding-domain is 5’-(A/G)CCC(T/G)CAT-3’or its compliment 5’-ATG(A/C)GGG(T/C)-3^12^. We found that this 8 bp nucleotide sequence was evolutionarily conserved in platypus and/or echidna orthologs of at least five *GCM1* target genes (Figure 5, Supplementary Data File 1). In order to further investigate the evolutionary conservation of GCM activity in the context of placenta development, we analysed GCM target gene orthologues in the chicken, which exhibits *GCM1* expression in the chorioallantoic membrane during embryogenesis^23^. Interestingly, GCM target motifs were also conserved in chicken orthologues of GCM target genes. This suggests that the regulatory network between *GCM1* and its target genes involved in early placental development or embryogenesis is ancient and predates the emergence of mammals. *GCM1* may therefore possess an ancient role in maternal-fetal nutrient exchange in addition to its role in syncytiotrophoblast formation in therian mammals.

Co-option of transposable elements has played a major role in shaping the evolution of the placenta in mammals^49^. In particular, in the therian lineage, the insertion of endogenous retroviruses (ERVs) is thought to have facilitated the evolution of complex, invasive placentae, owing to their functions in fusogenicity and immunosuppression^50^. The ERV-derived syncytin genes *ERVW-1* and *ERVFRD-1*, essential to syncytiotrophoblast formation and the evolution of invasive placentae, are regulated by *GCM1* in eutherian mammals^18,19^. We observed that the vast majority of the recently described putative syncytin genes in the echidna contain GCM DNA-binding sites, including Env-Tac1, which has demonstrated fusogenic activity^35^. Env-Tac1 is expressed in the kidney in echidnas, as is *GCM1*, suggesting a non-placental role for these genes in the echidna. Further work can explore the function of these genes in reproductive tissues in order to better understand their role in the non-invasive placentae of egg-laying species.

## Conclusion

The monotremes represent the most basal surviving mammalian lineage and their unique reproductive biology, including the development of a simple yolk sac placenta, provides a fascinating system in which to study the evolution of placentation. Our analysis of GCM genes in monotremes reveals the chromosomal relocation of *GCM1* after the divergence of monotremes and therian mammals, which may have played a part in the evolution of complex, invasive placentae. We report the first observation of *GCM1* expression in the monotreme reproductive tract as well as expression of *GCM2* in the platypus ovary, suggesting the adoption of a species-specific role. The regulatory network between *GCM1* and its placental target genes was found to be conserved across all mammalian lineages and to some extent in chicken. Together, this suggests an ancient role for *GCM1* in maternal-fetal exchange, predating the emergence of the placenta in mammals.

## Methods

### Fluorescence *in situ* hybridisation of BAC clones

BAC clones containing platypus Gcm1 (OABb-307B16) and Gcm2 (OABb-373O17) were obtained by screening a male platypus BAC library that were hybridised for overnight at 60.0°C with radioactively-labelled PCR products from the expression analysis study. Filters were washed with 2X SSC solution for 15 minutes at 60.0°C. Platypus male metaphase chromosome spreads were prepared from previously established fibroblast cell lines. The physical localisation of the BAC clones was determined using fluorescence *in situ* hybridisation on platypus male metaphase spread as described in Grutzner et al 2004^51^. Images were taken using a Zeiss AxioImager Z.1 epiflourescence microscope equipped with a CCD camera and Zeiss Axiovision software.

### Multiple sequence alignment and phylogenetic analysis

GCM1 and GCM2 amino acid sequences from *X. laevis*, *G. gallus*, *O. anatinus*, *T. aculeatus*, *M. domestica*, and *H. sapiens* were aligned using the ClustalO plugin^52^ (default settings) in Geneious Prime® 2023.2.1. Phylogenetic analysis involved creating an alignment of GCM amino acid sequences across 24 amniotes (plus *D. melanogaster* as an outgroup) using the ClustalO Geneious Prime® plugin (default settings). The alignment was used to construct a tree using W-IQ-Tree^53^ (default settings). Accession numbers for amino acid sequences used can be found in Supplementary Data File 1.

### RNA extraction and RT-PCR

Total RNA was extracted from frozen tissues stored at -80°C using TRIzol® (Invitrogen) for platypus and mouse tissues and NucleoZOL (Machery-Nagel™) for echidna tissues following the manufacturer’s instructions. RNA was re-suspended in nuclease-free water and stored at -80°C. g-DNA was removed using the DNA-*free*™ DNA Removal Kit (Invitrogen™) according to the manufacturer’s instructions.

cDNA conversion was performed on platypus and mouse RNA using the SuperScript™ III Reverse Transcriptase kit (Invitrogen™); cDNA conversion was performed on echidna RNA using the iScript™ cDNA Synthesis Kit (Bio-Rad). Synthesised cDNA was stored at -20°C.

25 µl reactions containing 5.0 µl of 5X Green GoTaq® reaction buffer (Promega), 0.5 µl 10mM dNTPs, 0.5 µl each 10 µM forward and reverse primer, 0.5 µl of locally made *Taq* DNA polymerase, 0.5 µl of cDNA, and 17.5 µl nuclease-free H2O were prepared for PCR amplification for each of the platypus and mouse samples. For the echidna samples, 20 µL reactions were prepared containing 4 µL 5x Phusion® HF Buffer, 0.4 µL10mM dNTPs, 1 each 10 µM forward and reverse primer, 0.2 µL Phusion® High-Fidelity DNA Polymerase, 1 µL cDNA, and 12.4 µL nuclease-free H2O. All primer sequences can be found in Supplementary Data File 1. PCR cycling conditions for platypus and mouse samples were as follows: initial denaturation at 96.0°C for 3 minutes, followed by 27 (*β-actin*) or 40 (*Gcm1* and *Gcm2*) cycles of denaturation at 96.0°C for 30 seconds, annealing at 54.0°C for 30 seconds and elongation at 72.0°C for 1 minute, with a final extension at 72.0°C for 7 minutes. PCR cycling conditions for echidna samples were as follows: initial denaturation at 98.0°C for 3 minutes, followed by 34 (*β-actin*), 38 (*Gcm1*), or 35 (*Gcm2*) cycles of denaturation at 96.0°C for 30 seconds, annealing at 58.0°C for 30 seconds and elongation at 72.0°C for 1 minute, with a final extension at 72.0°C for 7 minutes. PCR products were visualised on a 1.5% agarose gel stained with SYBR Safe DNA Gel Stain (Invitrogen™). PCR products were excised and the DNA extracted using QIAquick gel extraction Kit (Qiagen) for male platypus BAC library screening and RNA *in situ* hybridisation probes. All amplified PCR products underwent Sanger sequencing to confirm that they were the expected sequence.

### RNA *in situ* hybridisation

PCR products from RT-PCR were cloned with the pGEM®-T Easy Vector System (Promega). Sense and antisense digoxigenin-labelled riboprobes were generated from linearised plasmids. Platypus testis, ovary, and female reproductive tissue samples (AEEC permit R.CG.07.03 (F.G.), NPWS permit A193) were fixed in Bouin’s fixative for 2 hours, washed with PFA several times and stored in 70% ethanol until samples were embedded in paraffin. Embedded paraffin sections were cut into serial sections of 6 μm thickness. Section slides were deparaffinised, dehydrated and treated with 1.2 µg/µl Proteinase K for 30 minutes. Tissue section slides were hybridised overnight with digoxigenin-labelled riboprobes at 50°C. After hybridisation, section slides were washed in a series of SSC solutions as follows: 2X SSC, 1X SSC, 0.5X SSC and 0.1 XSSC for 15 minutes each at 50°C. An anti-digoxigenin alkaline phosphatase conjugated antibody (Roche) was used to detect the digoxigenin signal in a dark chamber at room temperature for 1 hour. Slides were stained with NBT and BCIP (Roche) and were placed in a humidified chamber in the dark for less than a day. Colour reaction was stopped with pH 8.0 1X TE buffer for 5 minutes at room temperature. Slides were then dried and mounted in Entellen® (ProSciTech).

### Target gene analysis

GCM target gene orthologues were identified in the chicken, platypus, echidna, opossum, and human using the NCBI Orthologs database (accession numbers for all genes can be found in Supplementary Data File 1). Each of these sequences, as well as the 1000 bp upstream of the transcription start site, was imported into Geneious for further analysis. A search for the GCM target motif sequence was performed, including the reverse complement in order to capture both DNA strands, and each motif was highlighted using the Geneious annotation tool. The annotated target gene orthologues were then aligned using the ClustalO Geneious plugin.

## Authors’ contributions

IW, DDMH and FG designed the study. IW and DDMH performed the experiments. FG provided samples. IW, DDMH and FG wrote the manuscript.

## Acknowledgements

This work was supported by the Australian Research Council (DP110105396). I.W. was supported by a University of Adelaide Research Scholarship. D.D.M.H. was supported by a postgraduate scholarship of the Ministry of Higher Education Malaysia and the University of Malaya.

