## Supplementary Figure 1 for "Evolution and expression of Glial Cells Missing (*GCM1* and *GCM2*) in monotremes suggests an ancient role in reproduction and placentation"

**Figure S1:** **Phylogenetic tree of GCM1 and GCM2 amino acid sequences from amphibians, reptiles, monotremes, marsupials, and eutherians.** Drosophila gcm was used as an outgroup. Sequences were aligned using ClustalO and consensus tree was constructed using IQ-TREE (both default settings). Node labels indicate bootstrap values.
